# Human hepatic tryptophan 2,3-dioxygenase ubiquitin-dependent protein degradation: The critical role of its exosite as the molecular lynchpin of its substrate-mediated protein stabilization

**DOI:** 10.1101/793380

**Authors:** Sung-Mi Kim, Yi Liu, YongQiang Wang, Shay Karkashon, Ariel Lewis-Ballester, Syun-Ru Yeh, Maria Almira Correia

## Abstract

Hepatic tryptophan 2,3-dioxygenase (TDO) is a cytoplasmic homotetrameric hemoprotein and the rate-limiting enzyme in the irreversible degradation of the essential amino acid *L*-tryptophan (*L-Trp*) to N-formylkynurenine, thus controlling the flux of *L*-Trp into its serotonergic and kynureninic/NAD pathways. TDO has long been recognized to be substrate-inducible via protein stabilization, but the molecular mechanism of this stabilization has remained elusive. Recent elucidation of human TDO (hTDO) crystal structure has identified a high-affinity (Kd ≈ 0.5 μM) Trp-binding exosite in each of its 4 monomeric subunits. Mutation of the Glu_105_, Trp_208_ and Arg_211_ comprising this exosite not only abolished the high-affinity *L*-Trp binding, but also accelerated the ubiquitin-dependent proteasomal degradation of hTDO. We have further characterized this hTDO degradation by documenting that its ubiquitination by gp78/AMFR and CHIP E2/E3 ligase complexes occurs on external Lys-residues within or vicinal to acidic Asp/Glu and phosphorylated pSer/pThr (DEpSpT)-clusters. Furthermore, we have identified the unstructured hTDO N- and C-termini as imparting relatively high proteolytic instability, as their deletion (ΔNC) markedly prolonged hTDO t_1/2_. Additionally, although previous studies reported that upon hepatic heme-depletion, the heme-free apoTDO turns over with a t_1/2_ ≈ 2.2 h relative to the t_1/2_ of 7.7 h of holoTDO, mutating the axial heme-ligating His_328_ to Ala has the opposite effect of prolonging hTDO t_1/2_. Most importantly, introducing the exosite mutation into the ΔNC-deleted or H328A-mutant completely abolished their prolonged half-lives irrespective of *L*-Trp presence or absence, thereby revealing that the exosite is the molecular lynchpin that defines *L*-Trp-mediated TDO induction via protein stabilization.

## INTRODUCTION

The hepatic hemoprotein tryptophan 2,3-dioxygenase (TDO; previously aka tryptophan pyrrolase) is the rate-limiting enzyme in the irreversible oxidative breakdown of the essential amino acid, *L*-tryptophan (*L*-Trp) to N-formylkynurenine (NFK) (1–7). This reaction is also carried out by indoleamine 2,3-dioxygenase (IDO) albeit with a much lesser substrate selectivity, as it also oxidizes many other indoles (4–6, 8). Unlike TDO, which is relatively selectively localized in liver and brain, IDO exhibits a much broader tissue distribution (4–6, 8). The product generated from either TDO- or IDO-mediated Trp/indole-breakdown is then further converted into various physiologically and pathologically relevant derivatives, including the cofactor NAD^+^ (4–8). Hepatic TDO is known to critically regulate the flux of *L*-Trp into serotonergic and kynureninic/NAD pathways, and is not only physiologically relevant in the control of serotonergic tone in the central and peripheral nervous systems (9–12), but also pathologically relevant in many human diseases (i.e. cancer, CNS depression, immune suppression, schizophrenia) (9–24). As such, along with IDO, it is currently a target of chemotherapeutic drug development (24–30).

The recently reported crystal structures of the human liver TDO (hTDO) holohemoprotein verifies that hTDO is indeed a homotetrameric hemoprotein and a dimer of dimers (*Fig. 1A*) (31). Each of the 4 monomeric hTDO-subunits is an all-helical structure with a 4-helical bundle arrangement constituting the tetrameric interface, and containing at one end a prosthetic heme moiety that utilizes His_328_ in the C-terminal end of helix J as its proximal ligand and an O_2_-molecule as its distal ligand (*Fig. 1B*). A *L*-Trp molecule is nestled above this heme-bound O_2_, strategically placed for the insertion of the O_2_-atoms into its indole C2-C3 double bond. This *L*-Trp is tethered by various residues including His_76_ (Helix B) with which it H-bonds (31). Site-directed mutation of His_76_ to Ser or Ala impairs the *k*_cat_ of the enzyme (32).

**Fig. 1.**
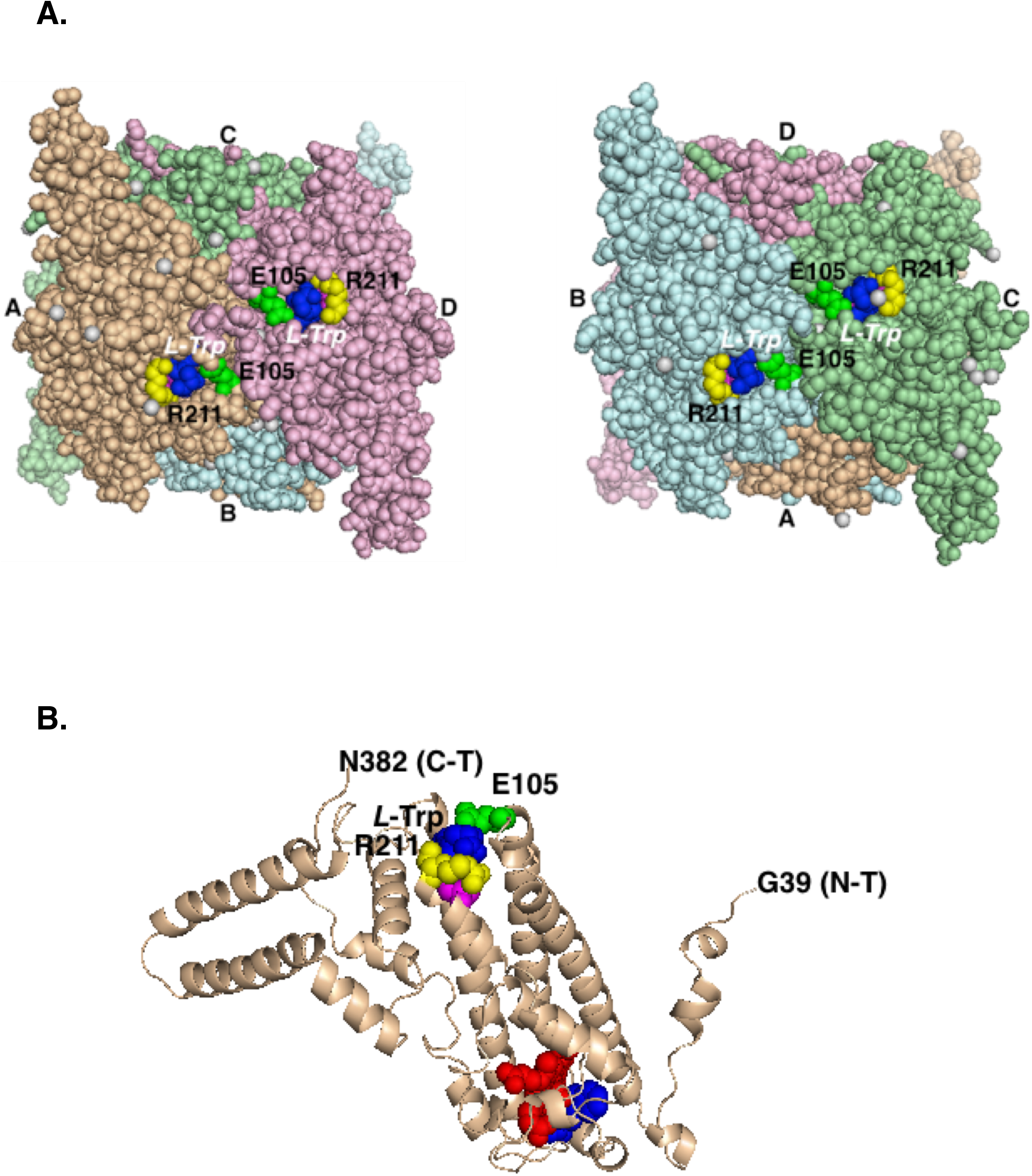
hTDO structure. **A.** Two opposite views of the homotetrameric crystal structure with each monomeric structure shown in different colors with a *L*-Trp molecule (deep blue) bound at each of the 4 exosites. Residues E_105_ and R_211_ of each exosite are depicted in green and yellow, respectively. **B**. Ribbon depiction of monomeric hTDO subunit B, an all-helical structure, curtailed at N-terminal residues 1-38, and C-terminal residues 383-4xx. At one end of the 4-helical bundle arrangement constituting the tetrameric interface, is the prosthetic heme moiety (shown as red spheres) that utilizes His_328_ in the C-terminal end of helix J as its proximal ligand and an O_2_-molecule as its distal ligand. A *L*-Trp molecule (deep blue) nestles above this heme-bound O_2_ at the active site. A second *L*-Trp molecule (deep blue) is shown bound to an “exosite” on hTDO’s external surface, at the other end of its 4-helical bundle, 42 Å away from the prosthetic heme. Residues W208, E_105_ and R_211_ of each exosite are depicted in magenta, green and yellow, respectively.

Although this crystal structure shows that all 4 active sites of the hTDO molecule are complexed with heme, the endogenous hepatic heme availability permits only 2 of the subunits to be physiologically saturated with heme at any given time, and the enzyme exhibits ≈ 50% of its full catalytic capacity (33, 34). Addition of heme *in vitro*, or injection of exogenous heme *in vivo* can restore hTDO to its full catalytic potential (33, 34). Physiologically, the enzyme is in equilibrium with the hepatic “free-heme pool” (33), thereby serving as a “hepatic heme-sensor”.

Early studies with purified *Pseudomonas* TDO have revealed that the enzyme exhibits sigmoidal kinetics at low *L-*Trp concentrations (35, 36). Inclusion of α-methyltryptophan (αMeTrp), an *L*-Trp derivative that cannot bind to the TDO active site due to steric clashes of its αMe-moiety with the prosthetic heme, converts its sigmoidal reaction kinetics at low [Trp] to hyperbolic, thereby revealing that the enzyme is allosterically activated (35, 36). Indeed, the recently reported hTDO crystal structure firmly verified that a second *L*-Trp molecule is bound on hTDO’s external surface to an “exosite”, at the other end of its 4-helical bundle, 42 Å away from the prosthetic heme (*Fig. 1B*) (31). At this exosite, *L*-Trp is sandwiched between Trp_208_ (helix H) and Pro_213_, and is in direct contact with several other residues, enabling bidentate ion-pair interactions of its main-chain carboxylate group with the side chain of Arg_211_ (helix H) and of its ammonium ion positioned near the main-chain carbonyl of Arg_103_ and the side chain of Glu_105_ (*Fig. 1B*) (31). Furthermore, isothermal calorimetric (ITC) analyses of hTDO revealed that the *L*-Trp affinity to this exosite (K_d_ ≈ 0.5 μM) is much higher than that of *L*-Trp at the active site (K_d_ ≈ 54 μM) (31). Site-directed mutagenesis of E_105_, W_208_ and R_211_ (hence named the EWR-or exosite mutant) to LVL abolished the high affinity *L*-Trp-binding, but retained its low-affinity *L*-Trp-binding (K_d_ ≈ 60 μM). However, this exosite hTDO mutant exhibited little alteration of its catalytic turnover (*k*_cat_) or K_m_ (31).

Classic studies of Schimke & coworkers (37–39) have documented the proteolytic stabilization of hepatic TDO by *L*-Trp-administration to rats and established the then radical concept of “*substrate-mediated enzyme induction via protein stabilization*” in mammals. Prompted by these earlier findings and our most recent structural discovery of an exosite binding of *L*-Trp, we examined whether this site was involved in substrate mediated hTDO protein stabilization (31). Indeed, relative to the wild type (WT) hTDO, its exosite EWR-mutant not only exhibited a much shorter half-life (t_1/2_), but unlike the WT-enzyme, its t_1/2_ was not prolonged in the presence of α-MeTrp, the analog capable of extending TDO t_1/2_ through binding to this exosite (31). This revealed that the TDO exosite was indeed relevant to its *L*-Trp-elicited protein stabilization. However, this site is apparently, not the only structural determinant of TDO-protein stability. Hepatic heme depletion and consequent deprivation of heme for prosthetic binding has also been found to accelerate its t_1/2_ from 7.7 h for the TDO holohemoprotein to 2.2 h for the apoprotein in rat livers (34). These t_1/2_s were extended to 11.4 h and 6.7 h, respectively, upon *L*-Trp (500 mg/kg, ip) administration to rats (34). Thus, the ability to bind prosthetic heme was a plausible TDO t_1/2_ determinant.

In this manuscript, we characterized the proteolytic degradation of hepatic hTDO by identifying the ubiquitin (Ub)-dependent proteasomal degradation (UPD) as its major degradation pathway, and the structural degrons and/or determinants relevant to its UPD. Our findings reveal that although hTDO possesses multiple structural features relevant to its protein stability, the exosite appears to be by far the major determinant of its UPD.

## Materials and Methods

### Expression Plasmids

Full-length (FL) hTDO expression plasmid (pcDNA6-hTDO-FL) was constructed by inserting a DNA fragment encoding the full-length human TDO into pcDNA6-(His)_6_ vector. To determine the role of the substrate binding site in TDO degradation based on our previous study (31), three hTDO amino acids (E_105_, W_208_, and R_211_) were mutated by QuickChange Lightning mutagenesis kit (Agilent Technologies, Santa Clara, CA, USA) to generate pcDNA6-hTDO-EWR mutant (E105L/W208V/R211L triple mutant). To determine the role of the heme-binding sites in TDO degradation and stabilization, His_328_ and His_76_ residues of full length-hTDO or EWR mutant-hTDO, were replaced with Ala using a site-directed mutagenesis kit, as described (40). N-terminal (Δ1-38), C-terminal (Δ389-406), and N-/C-terminal (ΔNC; Δ1-38/Δ389-406) deletion mutants of hTDO-FL or hTDO-EWR were also constructed. All constructs were verified through DNA sequencing analyses.

### Cell culture and treatment

Human hepatocellular HepG2 cells, were maintained at 37°C under 5% CO_2_ in Minimum Essential high glucose medium (MEM) containing nonessential and essential amino acids (EAA, including 0.050 mM L-Trp), 10% fetal bovine serum (FBS), 100 IU/ml penicillin, and 100 μg/ml streptomycin. For inhibition of hTDO degradation, HepG2 cells were transfected with pcDNA6-TDO-FL and treated 48 h later with the UPD inhibitor MG132 (20 μM), or autophagic lysosomal degradation (ALD) inhibitors 3-methyladenine (3-MA; 5 mM) and NH_4_Cl (10 mM).

### ^35^S-L-Met/Cys-pulse-chase analyses

HepG2 cells were transfected with the pcDNA6-hTDO-(His)_6_ or pcDNA6-EWR-(His)_6_ vector in 6-well-plates. After 48 h, culture medium was removed and replaced with methionine/cysteine-free MEM (containing 0.05 mM L-Trp along with other EAA), with or without added αMeTrp (2.5 mM) for 1 h, and subsequently pulsed with ^35^S-L-Met/Cys (75 μCi) for 1 h. After two washes with ice-cold PBS containing Met (0.2 mM)/Cys (1.4 mM), cells were further incubated with MEM containing cold Met (5 mM)/Cys (5 mM) with or without added αMeTrp (2.5 mM), for 0, 30, 60 and 90 min at 37 °C. The cells were lysed with Cell-Signaling Lysis buffer, containing general protease/phosphatase inhibitors, and N-ethylmaleimide (10 mM) to inhibit cellular deubiquitinases. The lysate was sedimented at 10,000 g at 4 °C for 10 min to remove insoluble cell debris, and its protein concentration was determined by the bicinchoninic assay (BCA). Lysate protein (200 μg) was then diluted (1:4, v:v) in Dynabead pull-down buffer and mixed with Dynabeads (50 μl), and incubated at 4 °C with rotation overnight. The Dynabeads-His_6_-tagged hTDO protein complexes were then collected using a magnetic stand and washed five times with Dynabead-washing buffer. The His6-tagged hTDO proteins were eluted by heating the complexes for 5 min in an SDS-PAGE sample-loading buffer [40 μl, 62.5 mM Tris buffer containing 25% (v/v) glycerol, 10% (w/v) SDS, 5% (v/v) β-mercaptoethanol and 0.01% (w/v) bromophenol blue]. The radioactivity of a 10 μl-aliquot was monitored in 4 ml of Ecolume using a Beckman LS3801 liquid scintillation counter, and another 30 μl-aliquot of the eluate (containing parent TDO protein and its ubiquitinated species) was subjected to SDS-PAGE (4–12%). The gels were dried, exposed for 2-3 days, subjected to fluorography with Typhoon scanning.

### Cycloheximide (CHX)-chase hTDO turnover analyses

We compared the relative stability of hTDO-His_6_ and its structural mutants by transfection of each His_6_-tagged construct for 48 h into HepG2 cells. CHX was then added to block *de novo* synthesis, and hTDO or mutant proteins pulled-down with Dynabeads at various times thereafter, and subjected to Western-immunoblotting with both anti-His6 and anti-TDO IgGs. Transfection of each mutant construct and CHX treatment were performed in 3-6 individual cell cultures to obtain 3-6 biological replicates. Western immunoblot images were quantified using ImageStudioLite (Licor Corp.) and normalized to the 0 min control for each individual experiment. The data were then analyzed by Prism Graphpad, and the half-life for each mutant was calculated on the basis of the nonlinear fit of exponential decay using data points from each of the biological replicates.

### hTDO Ubiquitination

This was determined exactly as detailed previously with the fully reconstituted gp78/AMFR and CHIP-E2/E3 ligase ubiquitination systems containing hTDO or its deletion or site-directed mutants as the substrate (31).

### hTDO Phosphorylation

This was monitored using each of the protein kinases singly or in combination in a phosphorylation system containing purified hTDO-FL as the substrate detailed previously (41, 42). hTDO-His_6_ was isolated with Dynabeads, subjected to trypsin/lysyl endoprotease C digestion, and phosphorylated peptide digest subjected to LC-Mass spectrometric (MS/MS) analyses and phosphorylated peptide identification as previously detailed (41, 42).

### Western Immunoblotting analyses of hTDO and its deletion or site-directed mutants

His_6_-tagged hTDO or its deletion or site-directed mutants were subjected to immunoblotting analyses using an anti-TDO and/or anti-His_6_-IgGs as detailed previously (31).

## RESULTS & DISCUSSION

### UPD is the major pathway in TDO proteolytic turnover

We have previously reported that hTDO incurs ubiquitination that targets it to UPD (31). To examine the relative contribution of UPD versus ALD to hTDO proteolytic turnover, we employed diagnostic inhibitors of these pathways as probes in HepG2 cells transfected with a hTDO-His_6_ plasmid for 48 h. Upon treatment with the proteasomal inhibitor MG132, hTDO was greatly stabilized in these cells both as the parent 47 kDa-species as well as its UPD-targeted K_48_-linked ubiquitinated species (*Fig. 2*). Some stabilization of this hTDO parent protein and its ALD-targeted K_63_-linked ubiquitinated species was also observed upon treatment of these transfected cells with ALD inhibitors: 3-methyladenine (3-MA), an inhibitor of autophagosomal formation, coupled with NH_4_Cl, an inhibitor of lysosomal proteases that functions by alkalinizing the intralysosomal pH. These findings revealed that although UPD is its major intracellular degradation pathway, hTDO also incurs appreciable degradation via ALD (*Fig. 2*). Although hepatic hTDO is a cytoplasmic enzyme, it is ubiquitinated largely by the endoplasmic reticulum (ER)-polytopic E3 Ub-ligase gp78 and the cytosolic E3 Ub-ligase, CHIP, as previously reported (31). We have previously excluded the E3 Ub-ligase Hrd1, a significant ERAD-contributor in yeast, as relevant to hTDO ubiquitination (31). Through HepG2 cell coexpression of hepatic hTDO with gp78, CHIP, HRD1 or FL-TEB4/MARCH IV, a mammalian homolog of yeast Doa10, an ER E3 Ub-ligase involved in the ubiquitination of both cytosolic and ER-proteins, we now find that although TEB4/MARCH IV was more effective than HRD1, it was considerably more sluggish than either gp78 and/or CHIP E3 ligases (*data not shown*).

**Fig. 2.**
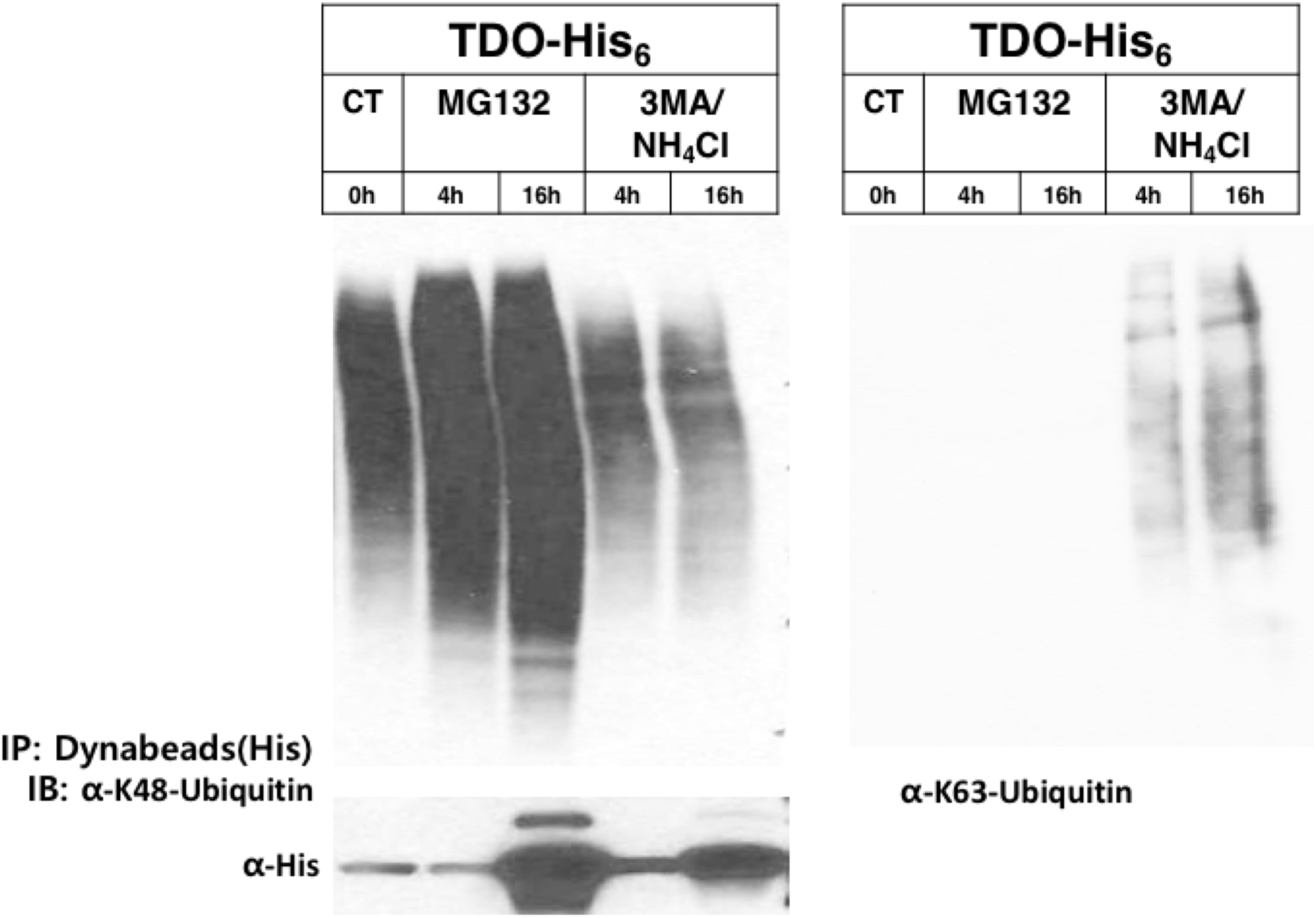
Relative hTDO turnover via UPD versus ALD. hTDO-His_6_ plasmid was transfected into HepG2 cells at 0 h and cells cultured for 48 h. Cells were then cultured with or without the proteasomal inhibitor MG132, or ALD inhibitors 3-MA/NH4Cl and harvested at 0, 4 and 16 h thereafter. Cell lysates were treated with magnetic Dynabeads to isolate the His_6_-tagged hTDO protein at each of these times. The isolated hTDO proteins were solubilized and subjected to Western immunoblotting against an anti-K_48_-linked or K_63_-linked Ub antibody or with anti-His_6_-antibody.

### gp78- and CHIP-mediated hTDO ubiquitination occurs within or vicinal to acidic DEpSpT-surface clusters

We have identified 13 sites of hTDO phosphorylated *in vitro* by various protein kinases using peptide mapping and LC-MS/MS analyses (Table 1) as previously detailed (41–43). These 13 sites along with the 15 previously identified Lys-residues ubiquitinated by both CHIP and gp78 E3 Ub-ligases (31) have been mapped on the recently reported hTDO homodimeric and homotetrameric structures (*Fig. 3*). Structural inspection of these ubiquitinated Lys-residues reveals that they not only lie largely on the external surface away from the hTDO dimeric interface, but also within or in close proximity to surface “DEpSpT” clusters of acidic Asp/Glu-residues and its phosphorylated Ser/Thr-residues. We have previously characterized through chemical cross-linking and LC-MS/MS analyses as well as site-directed mutagenesis, that such acidic, negatively charged residues either within disordered surface loops or clustered together by the tertiary protein fold were important for the molecular recognition of other gp78 and CHIP target substrates (i.e. human liver CYPs 3A4 and 2E1) by positively charged domains of these two E2/E3 Ub-ligase complexes (43). They thus essentially constituted phosphodegrons for molecular recognition of these E2/E3-substrates (43). The consistent finding of hTDO Lys-residues ubiquitinated by CHIP and gp78/AMFR within similar acidic surface DEpSpT-clusters reinforces the notion that such negatively charged clusters are indeed relevant for target substrate molecular recognition by these E3 Ub-ligases.

**Table 1.**
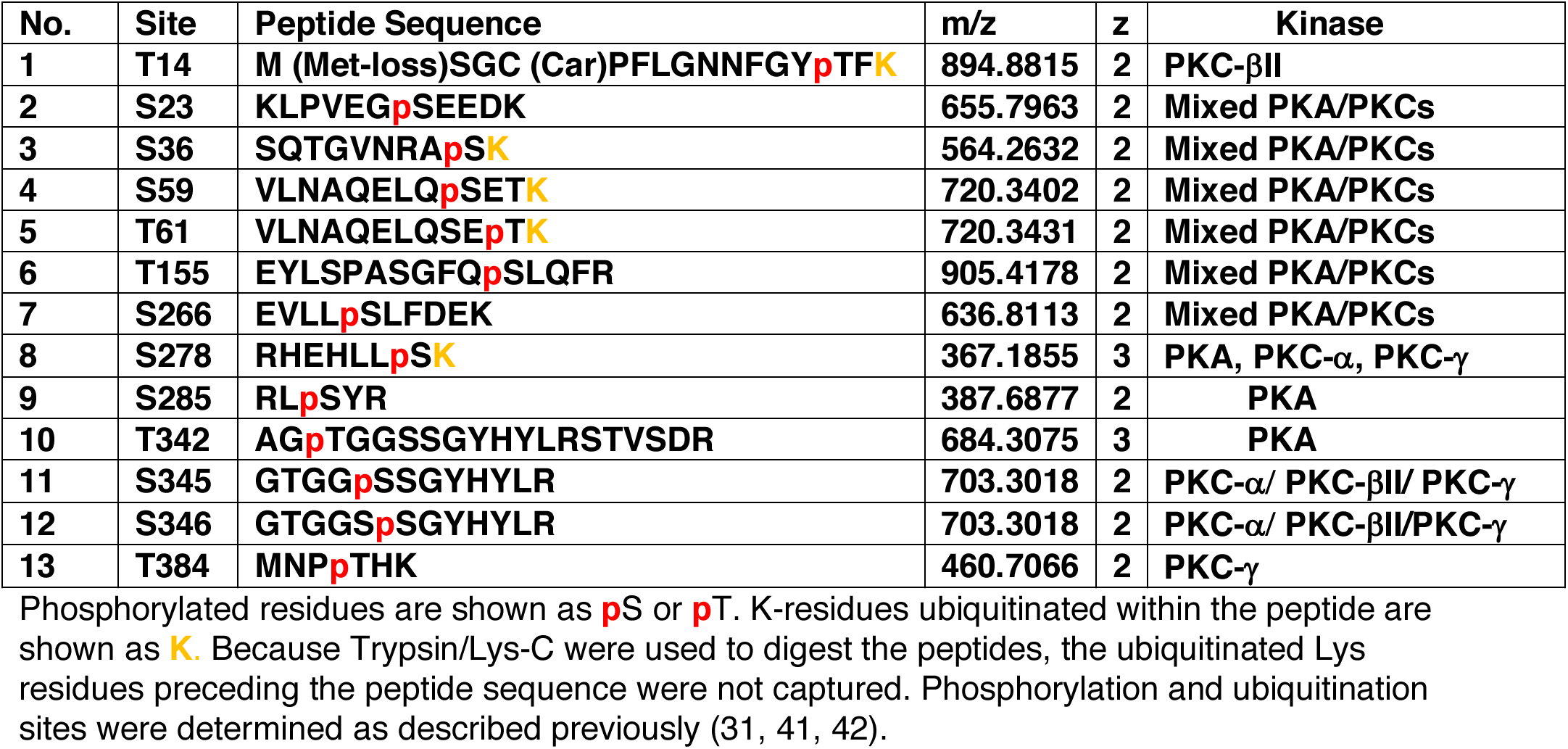
List of identified specific kinase-catalyzed TDO phosphorylation sites

| No. | Site | Peptide Sequence | m/z | z | Kinase |
| --- | --- | --- | --- | --- | --- |
| 1 | T14 | M (Met-loss)SGC (Car)PFLGNNFGYpTFK | 894.8815 | 2 | PKC-βII |
| 2 | S23 | KLPVEGpSEEDK | 655.7963 | 2 | Mixed PKA/PKCs |
| 3 | S36 | SQTGVNRApSK | 564.2632 | 2 | Mixed PKA/PKCs |
| 4 | S59 | VLNAQELQpSETK | 720.3402 | 2 | Mixed PKA/PKCs |
| 5 | T61 | VLNAQELQSEpTK | 720.3431 | 2 | Mixed PKA/PKCs |
| 6 | T155 | EYLSPASGFQpSLQFR | 905.4178 | 2 | Mixed PKA/PKCs |
| 7 | S266 | EVLLpSLFDEK | 636.8113 | 2 | Mixed PKA/PKCs |
| 8 | S278 | RHEHLLpSK | 367.1855 | 3 | PKA, PKC-α, PKC-γ |
| 9 | S285 | RLpSYR | 387.6877 | 2 | PKA |
| 10 | T342 | AGpTGGSSGYHYLRSTVSDR | 684.3075 | 3 | PKA |
| 11 | S345 | GTGGpSSGYHYLR | 703.3018 | 2 | PKC-α/ PKC-βII/ PKC-γ |
| 12 | S346 | GTGGSpSGYHYLR | 703.3018 | 2 | PKC-α/ PKC-βII/PKC-γ |
| 13 | T384 | MNPpTHK | 460.7066 | 2 | PKC-γ |

Phosphorylated residues are shown as pS or pT. K-residues ubiquitinated within the peptide are shown as K. Because Trypsin/Lys-C were used to digest the peptides, the ubiquitinated Lys residues preceding the peptide sequence were not captured. Phosphorylation and ubiquitination sites were determined as described previously (31, 41, 42).

**Fig. 3.**
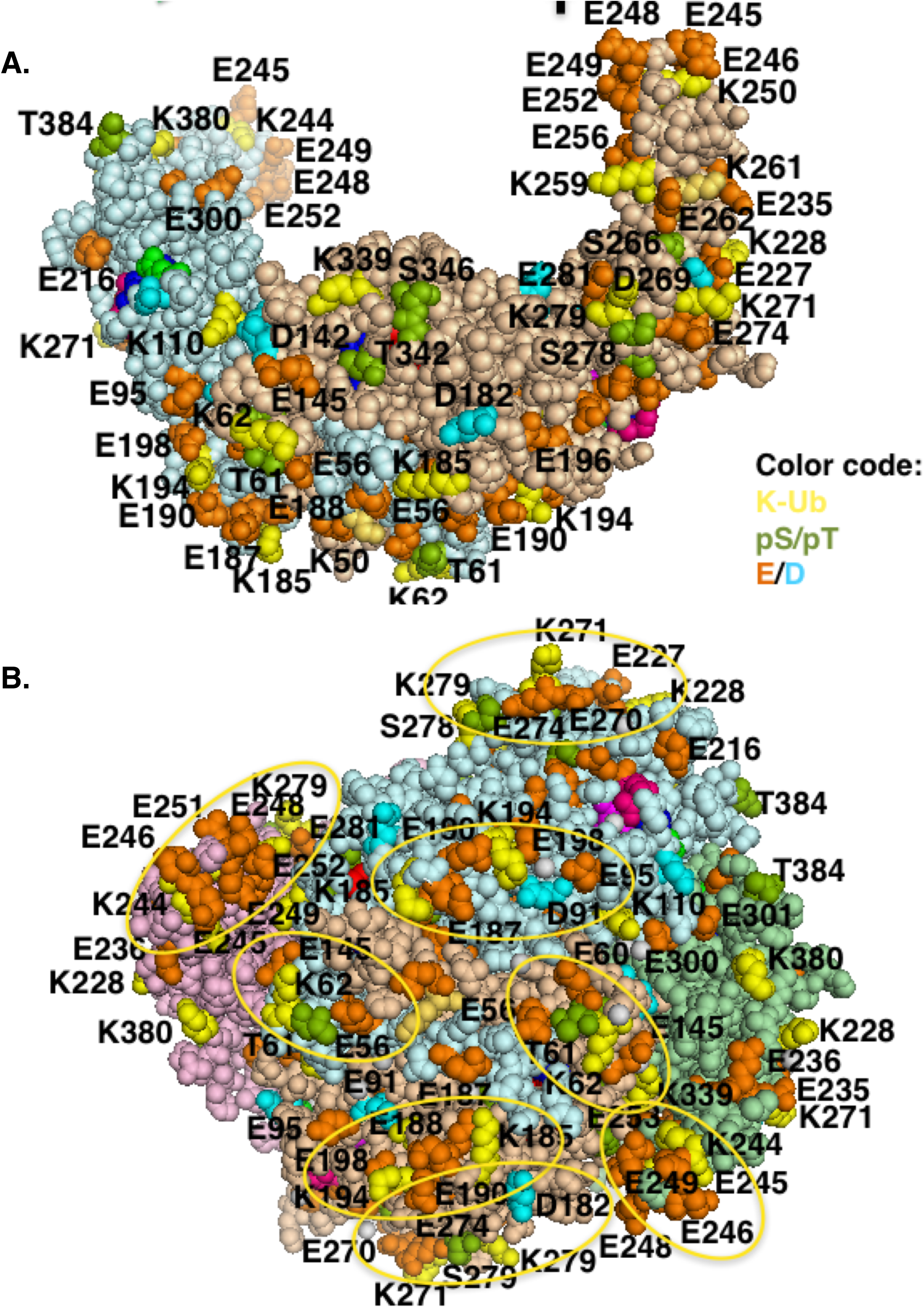
hTDO ubiquitinated K-residues vicinal to its surface DEpSpT-clusters. **A.** The surface of a hTDO dimer showing its ubiquitinated K-residues in yellow, its acidic D and E residues in cyan and orange, respectively, and the phosphorylated S/T sites identified through LC-MS/MS analyses in green. **B**. The hTDO tetrameric structure with the Ub-K residues vicinal to the DEpSpT clusters are encircled in yellow. Note that because the hTDO structure determined was of its ΔNC-tetramer, this depiction excludes the K and DEpSpT clusters in the unstructured regions of the deleted N- and C-termini.

### The relative importance of L-Trp-binding or the active site prosthetic heme-binding to hTDO protein stability

We have previously documented that the hTDO-exosite is very important for its α-MeTrp-mediated protein stabilization, as the site-directed mutation of key exosite EWR residues not only accelerated hTDO turnover, but also abolished its protein stability (31). Cycloheximide (CHX)-chase analyses reiterates these findings (*Fig. 4*). This TDO structural stability upon α-MeTrp/*L*-Trp-exosite-binding is imparted by the pulling together of the end of the 4-helical bundle distal to the active site heme and its helix-loop-helix domain (31). The structure of an α-MeTrp/*L*-Trp-free TDO protein surface is consequently much floppier with the exosite residues spread widely apart (44).

**Fig. 4.**
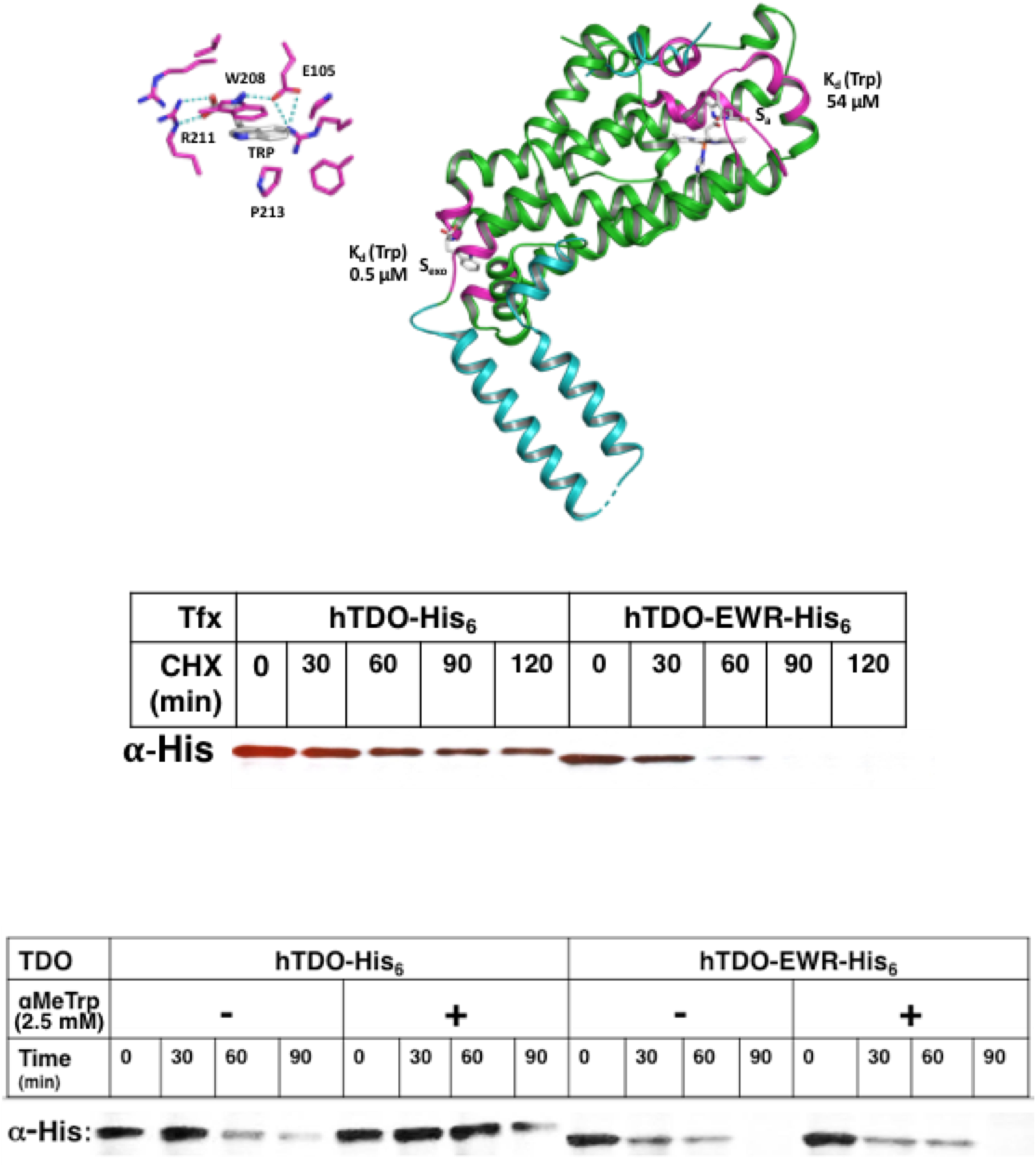
Relative effects of α-MeTrp on the turnover of hTDO and its EWR-exosite mutant. hTDO-His_6_ plasmid was transfected into HepG2 cells at 0 h and cells cultured for 48 h. CHX was added at this point, and cells were then cultured with or without α-MeTrp (2.5 mM) and harvested at 0, 30, 60, 90 and 120 min, thereafter. Cell lysates were treated with magnetic Dynabeads to isolate the His_6_-tagged hTDO protein at each of these times. The isolated hTDO proteins were solubilized and subjected to Western immunoblotting against an anti-His_6_-antibody.

However, as discussed above, it has long been recognized that another feature relevant to TDO protein stabilization is plausibly its prosthetic heme-binding, as the t_1/2_ of rat liver TDO holohemoprotein is accelerated upon depletion of hepatic heme (34). We previously have identified His_328_ and His_76_ through site-directed mutagenesis analyses of the 12 His-residues of a recombinant rat liver TDO as highly relevant to the expression of a spectrally detectable and functional TDO holohemoprotein (40). Indeed, the subsequent crystal structures of various TDO proteins have identified a His-residue corresponding to His_328_ as the proximal ligand for its prosthetic heme (31, 45, 46), whereas His_76_ was found relevant to *L*-Trp binding at the active site and subsequent catalytic turnover (32). For these reasons, we probed the role of prosthetic heme binding at His_328_ on hTDO proteolytic stability by CHX-chase analyses of a H328A-mutant transfected into HepG2 cells (*Fig. 5*). Surprisingly, we found that relative to the t_1/2_ (25.9 min) of the full-length WT hTDO or its EWR mutant (t_1/2_, 17.9 min), its H328A mutant was considerably more proteolytically stable with an even longer t_1/2_ of 105.8 min (*Fig. 5; Table 2*). These findings indicated that in the absence of hepatic heme depletion, disruption of the prosthetic heme axial H_328_-ligation failed to affect hTDO protein stability. This despite the fact that such H328A mutation disrupts hTDO hemoprotein structure and function as indicated by its impaired spectroscopic signature and Trp-dioxygenase activity, respectively (40). Thus, it is plausible that when heme is not depleted, heme, albeit not H_328_-axially ligated, persists sufficiently at the active site through hydrophobic interactions with the other active site residues so as to preclude gross perturbation of hTDO structural conformation that consequently triggers proteolytic degradation. However, coupling the H328A-mutation with the EWR-mutation significantly accelerated the t_1/2_ (26.3 min) of the double mutant (*Fig. 5*; *Table 2*), thereby underscoring the critical structural relevance of the *L*-Trp-exosite binding to protein-stability of the hTDO H328A mutant as well as the parent WT protein. The mutation of hTDO His_76_ to Ser tended to similarly stabilize the hTDO protein (*Table 2*). But coupling this mutation with the EWR mutation, once again considerably accelerated its degradation and reduced hTDO-H76S/EWR protein stability by almost 50% (*Table 2*).

**Table 2.** Half-life determination of TDO and its structural mutants via CHX-chase and ^35^S-pulse-chase analyses.

| **TDO-Protein** | **CHX-Chase**<br>**[t<sub>1/2</sub> (min)]** | **<sup>35</sup>S-Pulse-Chase**<br>**-αMeTrp** | **[t<sub>1/2</sub> (min)]**<br>**+αMeTrp** |
| --- | --- | --- | --- |
| **FL** | **25.9** | **90** | **172** |
| **EWR** | **17.9** | **60** | **68** |
| **H328A** | **105.8** |  |  |
| **H328A/EWR** | **26.3** |  |  |
| **ΔN** | **≥ 247.1** |  |  |
| **ΔC** | **≥ 247.1** | **252** | **360** |
| **ΔNC** | **247.1** | **210** | **275** |
| **ΔNC/EWR** | **33.7** | **60** | **70** |
| **C-DE->G** | **≥ 247.1** |  |  |
| **C-5S->G** | **≥ 247.1** |  |  |
| **H76S** |  | **230** |  |
| **H76S/EWR** |  | **118** |  |

The t<sub>1/2</sub>s were calculated using Prism Graphpad nonlinear fit of exponential decay of Mean ± SD data points obtained from at least 3 separately transfected cell cultures.
Note: The t<sub>1/2</sub>s obtained by <sup>35</sup>S-pulse-chase analyses are relatively longer than the corresponding ones obtained by CHX-chase analyses for the simple reason that the Met/Cys-deficient MEM contained 0.05 mM Trp required as an essential amino acid for optimal *de novo* protein synthesis, which was 100-fold higher than the exosite K<sub>D</sub> (0.5 μM). Nevertheless, the introduction of the EWR-mutation in every case resulted in a marked t<sub>1/2</sub> reduction of the TDO mutant. Whenever αMeTrp (2.5 mM) was included in the cultures, it could further prolong the t<sub>1/2</sub> of the TDO proteins that retained an intact exosite.

**Fig. 5.**
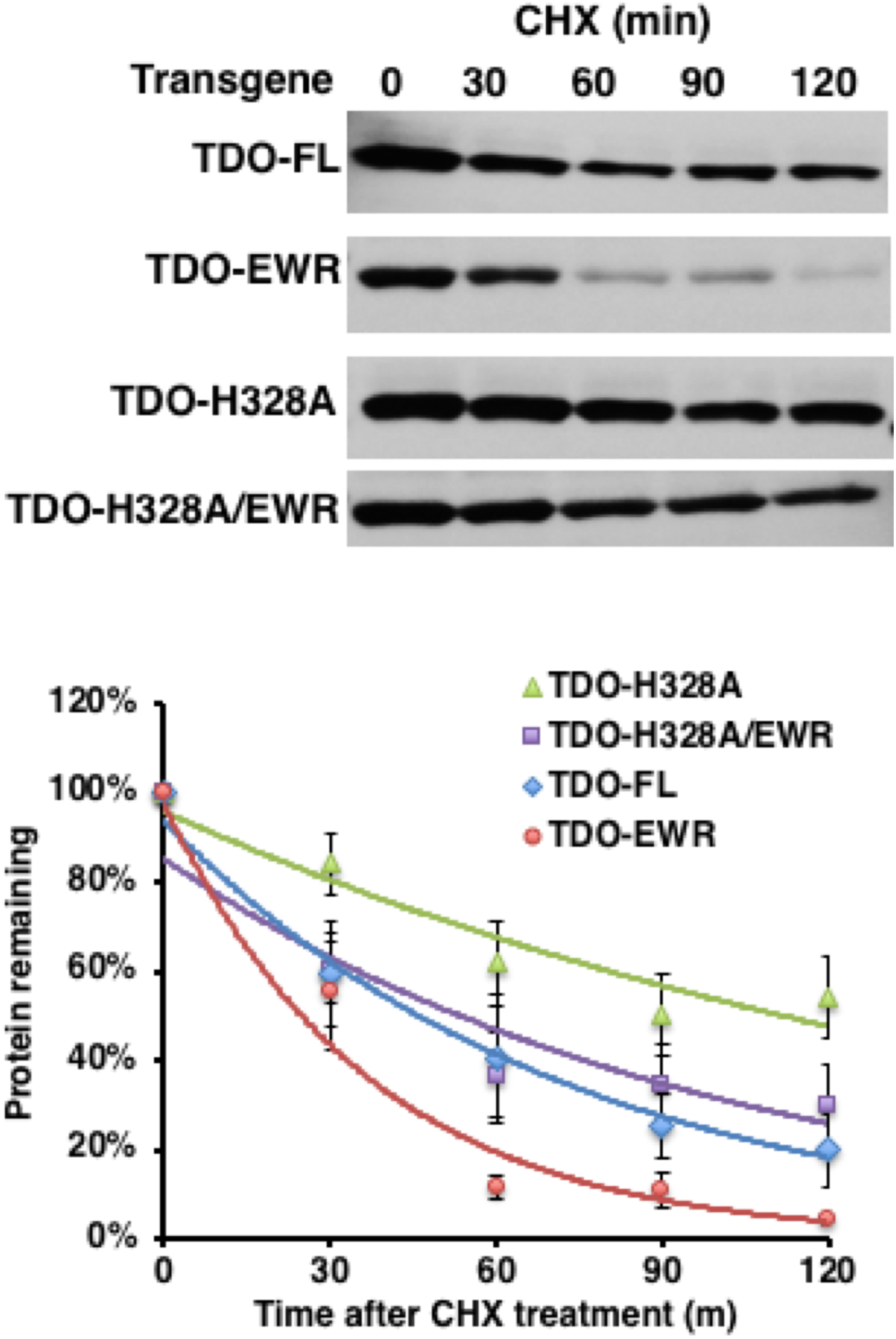
Relative turnover of hTDO, and its EWR-exosite, H328A and H328A/EWR mutants by CHX-chase analyses. For detailed methods see Materials & Methods. The top panel depicts a representative immunoblot out of at least 3 separate cell cultures. The bottom panel depicts the exponential decay using the Mean ± SD of those 3 individual experiments and Prism Graphpad software. The corresponding t_1/2_s are listed (Table 2).

### Relative importance of the unstructured N- and C-terminal domains to hTDO protein stability

Our past unsuccessful attempts at crystallization of the full-length recombinant rat liver TDO protein have led to the recognition that its relatively unstructured N- and C-termini interfere in the generation of crystals that could be successfully subjected to X-ray diffraction analyses. Indeed, deletion of these hTDO unstructured regions yielded diffractable crystals. Thus, our preliminary findings of HepG2 cell transfection of this N- and C-deleted (ΔNC) construct that had led to successful hTDO crystallization, revealed a highly stable protein (t_1/2_ 247.1 min) relative to the full-length WT hTDO, prompting us to interrogate the relative importance of both these terminal domains to physiologic hTDO protein stability. CHX-chase analyses indeed revealed that relative to WT hTDO, the N- or C-terminally deleted mutants were relatively stable (*Fig. 6; Table 2*). Coupling of the two deletions (ΔNC) also stabilized the protein relative to the full-length WT hTDO (*Figs. 6 & 7*). Indeed, this stability of the ΔNC-hTDO protein was associated with its tendency to be poorly ubiquitinated intracellularly relative to the WT hTDO in HepG2 cells (*Fig. 7, bottom panel*). Furthermore, in the presence of α-MeTrp, this tendency for poor ubiquitination could be further aggravated. This is consistent with the presence of a viable exosite conferring an even greater stability to the ΔNC-hTDO protein upon α-MeTrp-binding (*Fig. 7*). Disruption of the exosite on the other hand, markedly enhanced the ubiquitination and correspondingly reduced the stability of the ΔNC-hTDO protein, irrespective of the presence or absence of any α-MeTrp, thereby revealing that this exosite mutation overrides any stability conferred upon ΔNC-deletion (*Fig. 7; Table 2*). These findings were paralleled by those of ^35^S-pulse-chase analyses (*Table 2*). Accordingly, the t_1/2_ of the ΔNC-hTDO protein was markedly prolonged from t_1/2_ of 90 min for the WT-enzyme to a t_1/2_ of 210 min, and this was further prolonged to a t_1/2_ of 275 min upon αMeTrp-binding of the exosite of the ΔNC-hTDO protein. However, mutation of the exosite EWR-residues, markedly shortened the TDO t_1/2_ from 210 min to 60 min, and as previously, αMeTrp was incapable of appreciably extending the stability of this ΔNC-hTDO-EWR mutant protein (*Table 2*). Together these findings indicate that the disordered hTDO N- and C-termini impart relative proteolytic instability on the full-length protein, and their deletion substantially stabilized the protein. However, although this remarkable stability of the ΔNC-hTDO protein can be further extended upon αMeTrp-binding of its exosite, mutation of its key exosite EWR-residues override any such extended stability. Collectively, these findings once again underscore the major structural influence of the hTDO-exosite on its proteolytic stability.

**Fig. 6.**
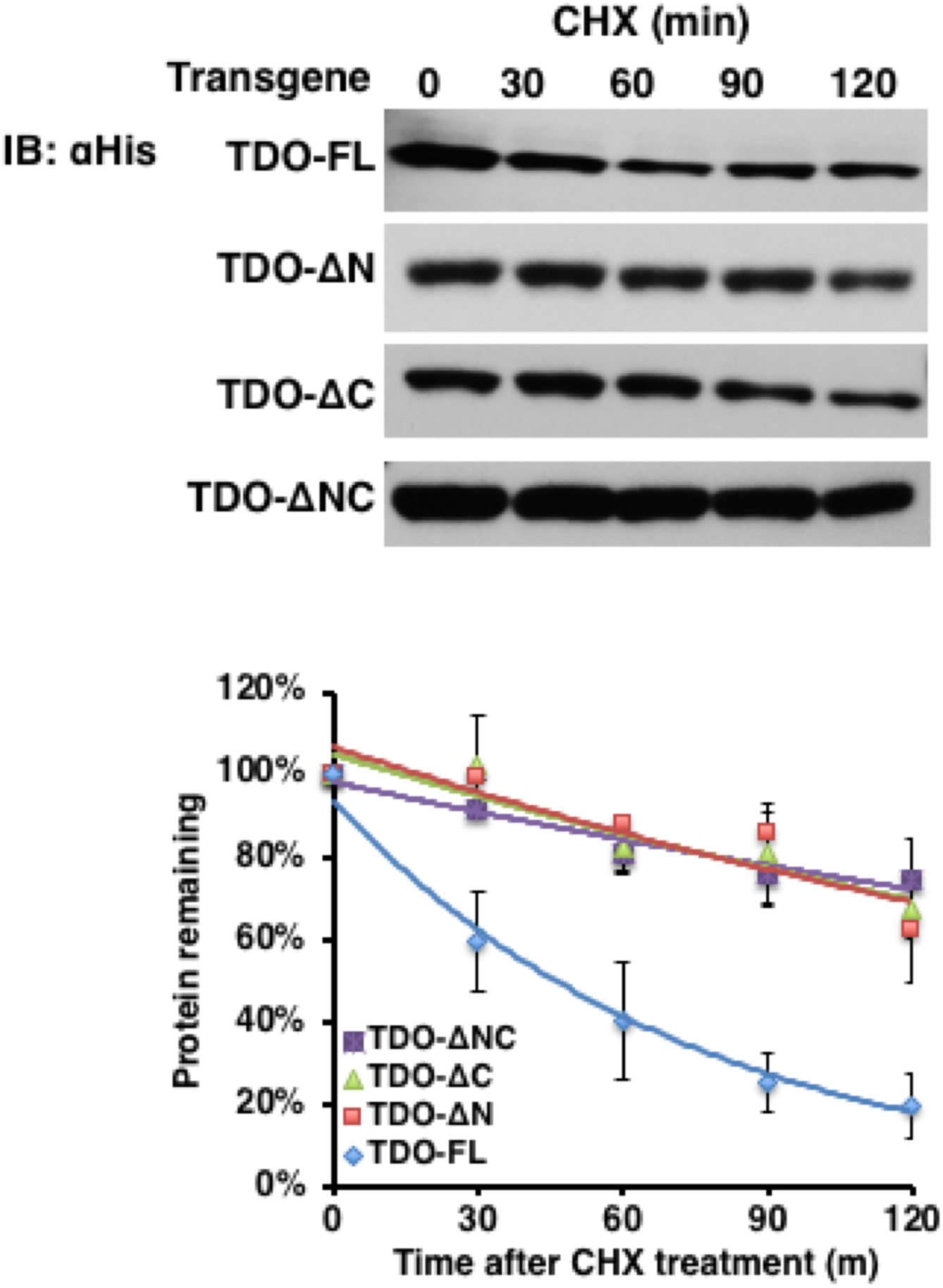
Relative turnover of full length (FL) hTDO, and its ΔNC-, ΔN- and ΔC-mutants by CHX-chase analyses. For detailed methods see Materials & Methods. The top panel depicts a representative immunoblot out of 3 separately cultured cells. The bottom panel depicts the exponential decay using the Mean ± SD of those 3 individual experiments and Prism Graphpad software. The corresponding t_1/2_s are listed (Table 2).

**Fig. 7.**
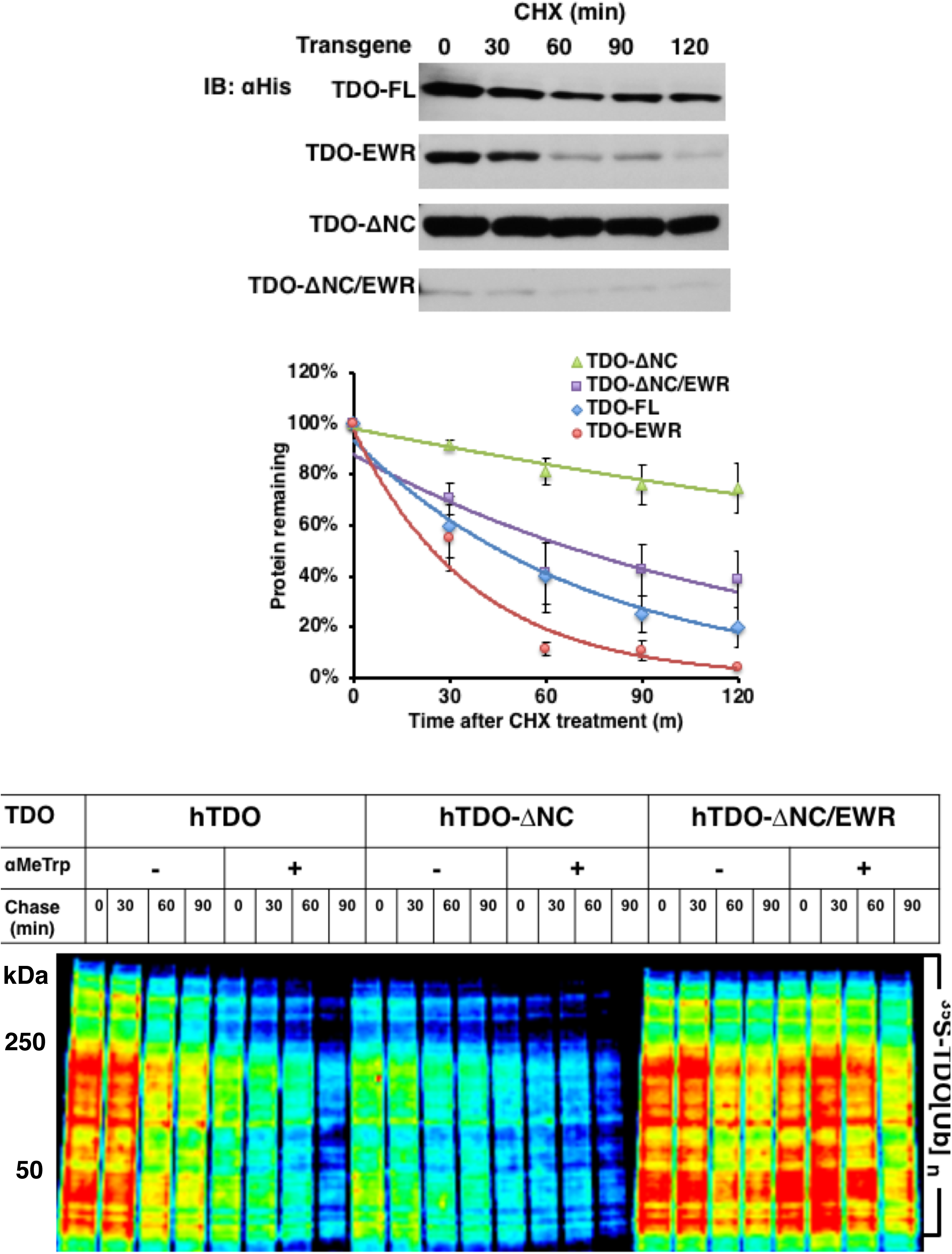
The influence of the EWR-exosite mutation on the proteolytic stability of the ΔNC-mutant, by CHX-chase analyses. **A.** For detailed methods see Materials & Methods. The top panel depicts a representative immunoblot out of 3 separately cultured cells. The bottom panel depicts the exponential decay using the Mean ± SD of those 3 individual experiments and Prism Graphpad software. The corresponding t_1/2_s are listed (Table 2). **B.** The relative ubiquitination of the FL, and its EWR-exosite, ΔNC- and ΔNC/EWR-mutants, in the presence or absence of α-MeTrp (2.5 mM). Note that α-MeTrp stabilizes the FL-protein against ubiquitination, whereas its stabilizing effect is greatly lost upon its EWR-mutant. On the other hand, the ΔNC-mutant is *per se* relatively stable to ubiquitination, and this stability is further enhanced by α-MeTrp. The color wheel intensity code is red>orange>yellow>green>blue>indigo>violet.

### Relative importance of certain C-terminal hTDO residues

Not only is the ΔNC-hTDO protein remarkably stable, but deletion of just the hTDO-C-terminus confers substantial proteolytic stability (*Fig. 6*). To identify the key C-terminal residues that may be relevant to its instability, we focused on the patch of acidic D/E-residues and the 5 potentially phosphorylatable S-residues. Mutation of D/E residues to Gly somewhat stabilized hTDO, relative to the WT-protein (*Fig. 8*). Similarly, mutation of the 5 S-residues also retarded hTDO degradation and stabilized the protein (*Fig. 8*). These findings indicate that the C-terminal D/E- and S-residues may play an active role in hTDO UPD. Although deletion of the first hTDO N-terminal 37 residues, stabilized the protein (*Fig. 6*), we have not similarly probed the role of the N-terminal K_17_EGSEEDKSQT_27_ subdomain in hTDO protein stability, although we have determined that the vicinal K_17_ and K_37_ residues in this N-terminal domain were indeed ubiquitinated (31). Nevertheless, we predict on the basis of the role of similar P450 surface DEpSpT-clusters in CYPs 3A4 and CYP2E1 (41–43) and other reported proteins (48, 49) that most likely, it will also be an important TDO structural feature.

**Fig. 8.**
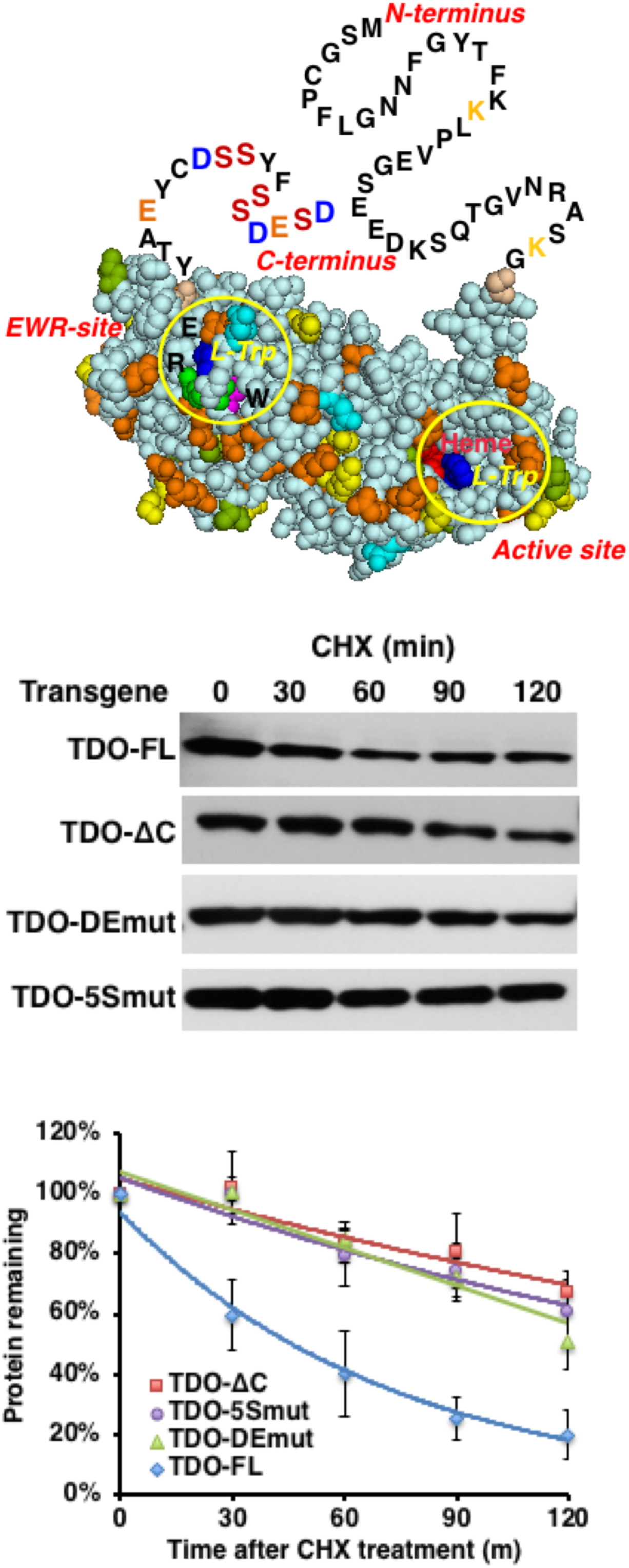
The influence of the 5 D/E-residues or 5S-residues in the unstructured hTDO C-terminus on its proteolytic stability. The top panel depicts a hTDO monomeric structure with its unstructured N- and C-termini included. For detailed methods see Materials & Methods. The middle panel depicts a representative immunoblot out of 3 separately cultured cells. The bottom panel depicts the exponential decay using the Mean ± SD of those 3 individual experiments and Prism Graphpad software. The corresponding t_1/2_s are listed (Table 2).

## Conclusions

The differences between glucocorticoid (GC)-mediated and substrate (*L*-Trp)-mediated protein induction have been long recognized (50–53). Employing *in vivo* isotope-labeling studies, Schimke et al. 1965 (38) first defined the underlying biological differences. Thus, while both physiological induction mechanisms enhance the hepatic TDO content, GCs such as cortisone and hydrocortisone increase the transcriptional rate of TDO synthesis via a GC-regulated element in the TDO promoter (53–55), whereas *L*-Trp does so by retarding the rate of TDO degradation (38). Hepatic TDO thus became one of the earliest classical examples of eukaryotic protein “*induction via protein stabilization*” (37–39), a biologic process until then only recognized in prokaryotes. However, the molecular basis for such *L*-Trp-mediated substrate stabilization remained elusive for nearly 50 years. The recently reported hTDO crystal structure has enabled us to elucidate the molecular mechanism of such *L*-Trp-mediated hTDO substrate stabilization, by identifying an “exosite” that not only bound *L*-Trp or αMeTrp with a very high affinity (Kd ≈ 0.5 μM), but which upon mutation of its critical circumscribing E_105_, W_208_ and R_211_ residues, greatly destabilized the hTDO protein structure and accelerated its proteolytic degradation via UPD (31). Herein, we show that indeed this “exosite” and not prosthetic heme axial ligation of hTDO is a dominant feature of its proteolytic stability. We also define other structural features conferring hTDO protein lability, such as its disordered N- and C-termini. Deletion of these two unstructured termini greatly enhanced hTDO proteolytic stability that enabled the successful crystallization and structural analyses of the hemoprotein. However, even this remarkable enhanced stability of the ΔNC-TDO deletion mutant is disrupted upon its exosite mutation. These findings thus underscore the critical structural relevance of this cardinal hTDO exosite feature as the molecular lynchpin of its *L*-Trp-mediated substrate-stabilization and hTDO regulation.

## Acknowledgments

We thank Prof. Mark Hochstrasser, Yale Univ, for a FL-TEB4/MARCH IV expression plasmid. We also acknowledge the use of the UCSF Bio-Organic Biomedical Mass Spectrometry Resource (Prof. A. L. Burlingame, Director) supported by the Adelson Medical Research Foundation.

